# Recovery of glutamatergic and GABAergic protein expression in visual cortex after monocular deprivation

**DOI:** 10.1101/684191

**Authors:** Justin L. Balsor, David G. Jones, Kathryn M. Murphy

**Author notes:** Correspondence should be addressed to Kathryn M. Murphy. Kathryn Murphy, Department of Psychology, Neuroscience & Behavior, McMaster University, McMaster University, 1280 Main Street West, Hamilton, ON L8S 4K1, Canada. Authors’ contributions: JB Designed research, performed research, analyzed data, wrote/revised the paper; DJ analyzed data, revised the paper; KM designed research, performed research, analyzed data, wrote/revised the paper.

## Abstract

A collection of glutamatergic and GABAergic proteins participate in regulating experience-dependent plasticity in the visual cortex (V1). Many studies have characterized changes to those proteins caused by monocular deprivation (MD) during the critical period (CP), but less is known about changes that occur when MD stops. We measured the effects of 3 types of visual experience after MD (n=24, 10 male and 14 female); reverse occlusion (RO), binocular deprivation (BD), or binocular vision, on the expression of synaptic proteins in V1 including glutamatergic and GABAergic receptor subunits. Synapsin expression was increased by RO but not affected by the other treatments. BD shifted the balance between glutamatergic and GABAergic receptor subunits to favor GABA_A_α1. In contrast, BV shifted expression to favor the glutamatergic mechanisms by increasing NMDAR and decreasing GABA_A_α1 subunits. None of the conditions returned normal expression levels to all of the proteins, but BV was the closest.

## Introduction

Glutamatergic and GABAergic proteins in visual cortex (V1) play central roles in regulating visual experience-dependent plasticity during the critical period (CP) (Hensch, 2005; Maffei and Turrigiano, 2008; Yashiro and Philpot, 2008; Smith et al., 2009; Heimel et al., 2011; Cooper and Bear, 2012; Hensch and Quinlan, 2018). Importantly, the subunit composition of those receptors can be changed by monocular deprivation (MD) including the delayed shift to more GluN2A-containing NMDARs (Fagiolini et al., 2003; Beston et al., 2010), and accelerated expression of GABA_A_α1-containing GABA_A_Rs (Fagiolini et al., 2004; Beston et al., 2010). Also, silencing activity in cultured visual cortex neurons increases GluA2-containing AMPAR at synapses (Gainey et al., 2009). Those MD induced receptor changes are linked with acuity deficits (Iny et al., 2006; Williams et al., 2015) suggesting that recovery of specific receptor proteins may support visual recovery.

Reverse occlusion (RO) is used to promote recovery by giving the deprived eye a competitive advantage, but the recovered acuity is transient and can be lost within hours of introducing binocular vision (Mitchell et al., 1984; Murphy and Mitchell, 1986; 1987). Binocular deprivation therapy (BD) has also been tried, but it too leads to poor acuity in both eyes (Duffy et al., 2015). In contrast, just opening the deprived eye and giving binocular vision (BV) supports long-lasting visual recovery in both eyes (Williams et al., 2015).

We measured the expression of glutamatergic and GABAergic receptor subunits in V1 of animals given either BV, RO or BD after MD. We compared subunit expression among the treatment 3 groups and with animals that had either normal binocular vision or MD.

## Materials & Methods

### Animals & Rearing Conditions

All experimental procedures were approved by the McMaster University Animal Research Ethics Board. We quantified the expression of 7 glutamatergic and GABAergic synaptic proteins in V1 of cats reared with MD from eye opening until 5 weeks of age and then given one of 3 treatments: RO for 18d, BD for 4d, or BV (1hr, 6hrs, 1d, 2d or 4d) (n=7, 4 male and 3 female). These rearing conditions are the same as described in Balsor et al., 2019 (see Figure 1 in Balsor et al., 2019) and the same dataset was used by that study. In addition, developmental data collected previously (Beston et al., 2010) from normal (2, 3, 4, 5, 6, 8, 12, 16, or 32 wks of age, n=9, 2 male and 7 female), or MDed animals (4, 5, 6, 9, or 32 wks, n=8, 4 male and 4 female) were used for comparison.

**Figure 1.**
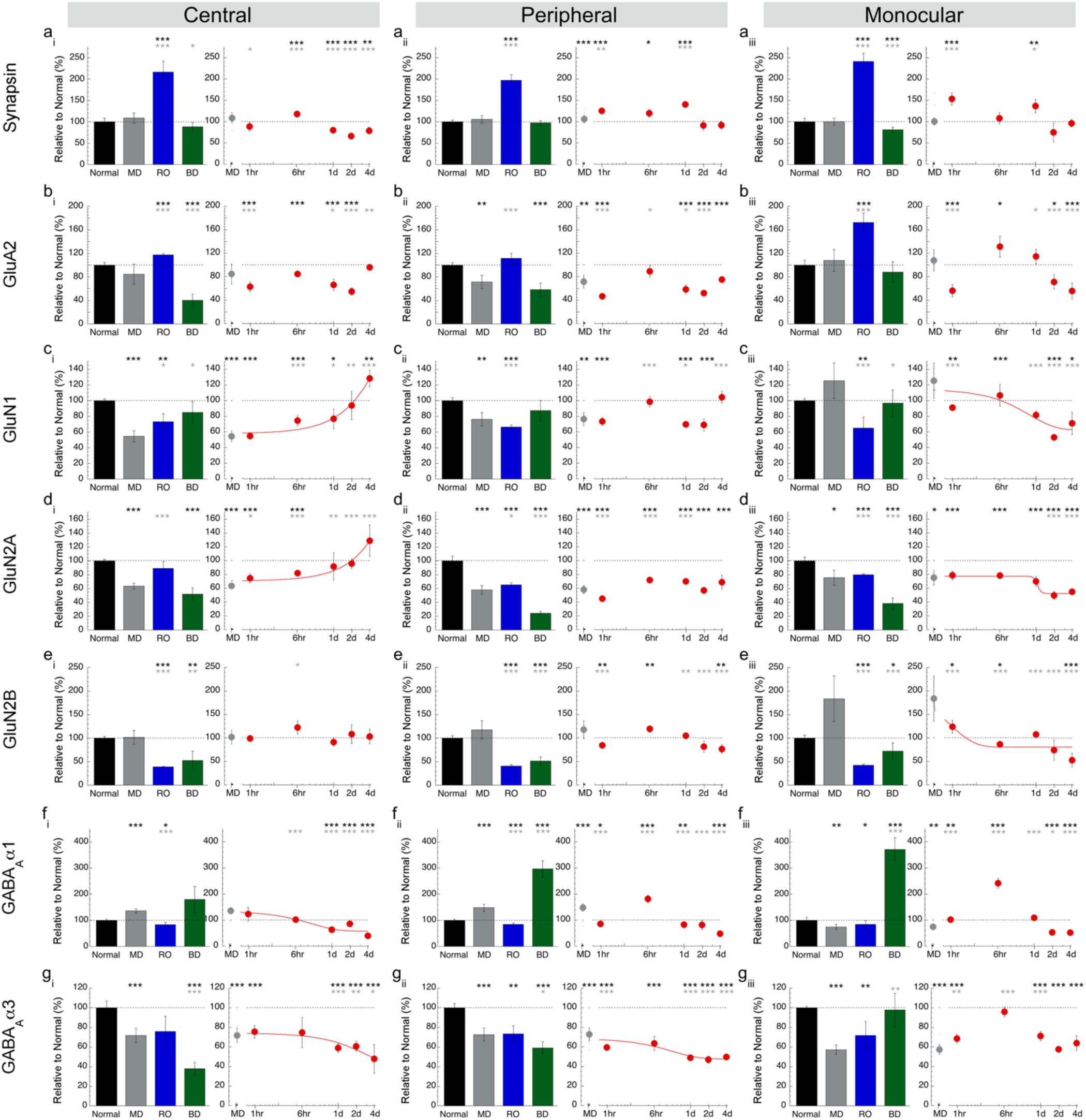
Expression of synaptic proteins in the different regions of visual cortex. Histograms showing the average expression relative to 5 wk normal animals for the 7 proteins (rows) and 3 regions of V1 (columns), normal 5 wk animals (black bars), animals reared with MD to 5wks (grey bars), and animals treated with either RO (blue bars) or BD (green bars). Scatter plots showing the average protein expression (red dots) after 1hr to 4d of BV treatment. When the trajectory of protein changes during BV treatment was well-defined by a function, curve fits were applied (red line). Error bars represent standard error of the mean (SEM). Black asterisks represent significant differences relative to 5wk normal, and grey asterisks represent significant differences relative to 5wk MD (*p<0.05,**p<0.01, ***p<0.001). The dotted black line on each graph represents 5wk normal expression. For exact p-values, Pearson’s R, and equations for the curve-fits see Table 3-1.

The procedures for MD, euthanizing and tissue collection were the same as described in Balsor et al., 2019.

### Synaptoneurosome preparation and Immunoblotting

Synaptoneurosomes were prepared, the protein concentrations were equated and immunoblotting was done according to the protocols described previously (Beston et al., 2010).

### Analysis of Protein Expression

The bands were identified on the autoradiographic film based on molecular weight and quantified using densitometry follow the protocol described previously (Beston et al., 2010).

The data were normalized relative to the average expression of the 5wk normal cases and plotted either as histograms to compare expression levels among the 5wk Normal, 5wk MD, RO, and BD animals or as scatterplots to follow expression changes over the 5 different lengths of BV. Table 2 summarizes the data.

**Table1:** List of primary antibody concentrations.

| Antibody | Concentration | Company | Lot Number | Location | RRID |
| --- | --- | --- | --- | --- | --- |
| anti-GluN1 | 1:2000 | BD Biosciences<br>Pharmingen | 556308 | San Diego, CA | RRID:AB_396353 |
| anti-GluN2A | 1:2000 | Millipore Sigma | 24826 | Burlington, MA | RRID: AB_95169 |
| anti-GluN2B | 1:2000 | Millipore Sigma | 28629 | Burlington, MA | RRID: AB_2112925 |
| anti-GluA2 | 1:1000 | Thermo Fisher |  | Waltham, MA | RRID: AB_2533058 |
| anti-GABAα1 | 1:500 | Santa Cruz<br>Biotechnology | L3102 | Santa Cruz, CA | |
| anti-GABAα3 | 1:2000 | Millipore Sigma | | Burlington, MA | |
| anti-Synapsin | 1:2000 | Thermo Fisher |  | Waltham, MA |  |

**Table 2:** The number of animals, cortical tissue pieces, and WB measurements for each condition and V1 region. Rows summarize the number of runs from the Central (C), Peripheral (P), and Monocular (M) regions of V1 within a rearing condition. The columns list each of the 7 proteins analyzed using Western blotting. Column sums detail the number of runs across rearing conditions and cortical areas. The information for Normal animals is in Table 2-1, and for MD animals is in Table 2-2.

| Number of Western Blot measurements after 2 replications |  |  |  |  |  |  |  |  |
| --- | --- | --- | --- | --- | --- | --- | --- | --- |
| Condition | Region | GluN1 | GluN2A | GluN2B | GABAα1 | GABAα3 | GluA2 | Synapsin |
| 5wk Normal | C | 4 | 4 | 4 | 4 | 4 | 4 | 4 |
|  | P | 16 | 16 | 16 | 15 | 16 | 16 | 16 |
|  | M | 4 | 4 | 4 | 4 | 4 | 4 | 4 |
| 5wk MD | C | 6 | 6 | 6 | 6 | 6 | 6 | 4 |
|  | P | 18 | 18 | 18 | 18 | 18 | 18 | 12 |
|  | M | 5 | 5 | 5 | 5 | 5 | 5 | 4 |
| 18d RO | C | 4 | 4 | 4 | 4 | 4 | 4 | 4 |
|  | P | 19 | 19 | 19 | 19 | 19 | 14 | 14 |
|  | M | 3 | 3 | 3 | 3 | 3 | 2 | 2 |
| 4d BD | C | 6 | 6 | 5 | 6 | 5 | 6 | 5 |
|  | P | 18 | 18 | 17 | 16 | 18 | 18 | 17 |
|  | M | 4 | 4 | 3 | 4 | 4 | 4 | 3 |
| 1hr BV | C | 4 | 4 | 4 | 4 | 4 | 4 | 4 |
|  | P | 16 | 16 | 16 | 16 | 16 | 16 | 16 |
|  | M | 4 | 4 | 4 | 4 | 4 | 4 | 3 |
| 6hr BV | C | 4 | 4 | 4 | 4 | 4 | 4 | 4 |
|  | P | 16 | 16 | 16 | 16 | 16 | 16 | 16 |
|  | M | 4 | 4 | 4 | 4 | 4 | 4 | 4 |
| 1d BV | C | 4 | 2 | 4 | 4 | 4 | 4 | 4 |
|  | P | 16 | 13 | 16 | 16 | 16 | 16 | 16 |
|  | M | 4 | 4 | 4 | 4 | 4 | 4 | 4 |
| 2d BV | C | 4 | 4 | 4 | 3 | 4 | 4 | 2 |
|  | P | 15 | 15 | 15 | 15 | 14 | 15 | 12 |
|  | M | 4 | 4 | 4 | 4 | 4 | 4 | 4 |
| 4d BV | C | 4 | 4 | 4 | 4 | 4 | 4 | 4 |
|  | P | 12 | 12 | 12 | 12 | 12 | 12 | 12 |
|  | M | 4 | 4 | 4 | 4 | 4 | 4 | 4 |
| SUM |  | 222 | 217 | 219 | 218 | 220 | 216 | 198 |

We analyzed heterogeneity in protein expression within a group by calculating an index of dispersion, the variance-to-mean ratio (VMR), for each protein and rearing condition. Proteins with VMR <1 were classified as under-dispersed, VMR=1 randomly dispersed, and VMR >1 were over-dispersed. We used this measure to compare among groups and assess if a rearing condition changed the variability of protein expression to make the group more or less heterogeneous.

### Statistical analyses

The mean and SEM for each protein and plasticity feature from the 3 regions of V1 was calculated for each rearing condition. We used the bootstrap resampling method because it is a conservative approach to analyzing small sample sizes when standard parametric or non-parametric statistical tests are not appropriate. Here bootstrapping was used to estimate the confidence intervals (CI) for each of the recovery groups and a Monte Carlo simulation was run to determine if the 5wk Normal or 5wk MD groups fell outside those CIs. The statistical software package R was used to simulate normal distributions with 1,000,000 points using the mean and standard deviation from the recovery groups (RO, BD, BV). Next, a Monte Carlo simulation randomly sampled with replacement from the simulated distribution *n* times, where *n* was the number of observations made from the normal or MD group (e.g., n = 4-19). The resampling procedure was repeated 100,000 times to determine the 95%, 99% and 99.9% CIs. The recovery group was considered significantly different (e.g., p<0.05, p<0.01 or p<0.001) from the normal and MD group if the mean of those groups fell outside the CI for the recovery group.

We tested if recovery during BV followed either an exponential decay or sigmoidal pattern by fitting curves to the data using Kaleidagraph (Synergy Software, Reading PA). Significant curve fits were plotted on the graphs to describe the trajectory of recovery.

## Results

### Analyzing recovery of synaptic proteins: Synapsin

We began analyzing the effects of the 3 recovery treatments by comparing expression of a pre-synaptic marker, synapsin, in the central, peripheral, and monocular regions of cat V1. As we reported previously(Beston et al., 2010), 5wks MD did not affect expression of synapsin relative to normals, but RO caused synapsin expression to double in all regions of V1 (C: 216%±25%, p<0.0001; P: 197%±13%, p<0.0001; M: 241%±19%, p<0.0001)(Figure 1a). In contrast, there was only slightly less than normal synapsin expression after BD (M: 82%±6%, p<0.0001) and 4d-BV (C: 79%±8%, p<0.0029).

### Analyzing recovery of synaptic proteins: Glutamatergic receptor subunits

Next, we quantified changes in GluA2 and GluN1 expression in V1. Here we found that RO promoted a small increase in GluA2 expression (C: 112%±8%, p<0.0001; P: 118%±2%, p=0.0691; M: 172%±16%, p<0.0001) but a loss of GluN1 (C: 73%±10%, p=0.0041; P:67%±3%, p<0.0001; M: 65%±14%, p=0.0019, Figure 1c). In contrast, after BD treatment GluA2 was reduced in the binocular regions of V1 (C: 40%±10%, p<0.0001; P: 58%±11%, p<0.0001).

BV treatment had variable effects on GluA2 similar to the timing of changes seen when short-term (hours) vs long-term (days) MD is started at the peak of the CP (Williams et al., 2015). GluA2 expression fluctuated during BV but tended to be slightly below normal levels (Figure 1b). Interestingly, the timing of the fluctuations was the same but in the opposite direction to what we found previously with 1d of MD (Williams et al., 2015). After 1d MD there was a decrease but here 1d BV had a transient increase in GluA2.

BV treatment had region-specific changes in GluN1 expression with central recovery to above normal levels (y = 297.43-239.58*exp(-x/11.61), df=25, R^2^=0.618, p<0.0001) but below normal levels in the monocular region (71%±14%, p=0.0214).

Next, we analyzed expression of the NMDAR subunits, GluN2A and GluN2B (Figure 3d,e), because they regulate ocular dominance and bidirectional synaptic plasticity in V1(Philpot et al., 2001; Cho et al., 2009) as well as receptor kinetics(Sun et al., 2017). Similar to our previous study(Beston et al., 2010), GluN2A expression was reduced after MD across all of V1 (C: 63%±4%, p<0.0001; P: 58%±6%, p<0.0001; M: 76%±11% =0.0220). RO promoted modest recovery of GluN2A in the central region (89%±10%, p=0.1367) but BD treatment did not. Instead after BD there was even less GluN2A than MDs in the peripheral (24%±2%, p<0.0001) and monocular regions (38%±8%, p<0.0001). BV promoted recovery of GluN2A in the central region (4d BV: 129%±23%, p=0.1022), but not in the other regions.

GluN2B expression was similar to normals after MD (Beston et al., 2010) but declined after both RO (C: 39%±1%, p<0.0001; P: 41%±3%, p<0.0001; M: 43%±2%, p<0.0001) and BD (C: 53%±20%, p=0.0085; P: 52%±8%, p<0.0001; M: 72%±18%, p=0.0354). BV, however, had little effect on GluN2B expression and after 4d it was similar to normal in the central region, and modestly reduced in peripheral (4dBV:77%±9%, p=0.0024) and monocular regions (4dBV: 53%±15%, p=0.0007).

### Analyzing recovery of synaptic proteins: GABAergic receptor subunits

Previous studies have shown that GABA_A_Rs are necessary for opening the CP(Hensch et al., 1998), that GABA_A_α1 regulates patterns of activity needed for ocular dominance plasticity(Fagiolini et al., 2004) and that GABA_A_α3 and GABA_A_α1 regulate the kinetics of GABA_A_Rs(Gingrich et al., 1995; Eyre et al., 2012). For these reasons, we examined how the treatments affected the expression of GABA_A_α1 and GAB_A_α3. We found that BD increased GABA_A_α1 expression (P: 297%±31%, p<0.0001; M:372%±45%, p<0.0001) but RO caused a slight decrease (C: 83%±8%, p=0.0239; P:85%±4%, p=0.0008; M: 85%±14%, p=0.0026). After 4d of BV GABA_A_α1 expression was below normal levels (4d BV C: 40%±9%, p<0.0001; P:48%±8%, p<0.0001; M: 51%±4%, p<0.0001). There was, however, a transient increase in GABA_A_α1 outside the central region after 6hrs of BV (P: 181%±14%, p<0.0001; M:242%±19%, p<0.0001). GABA_A_α3 expression remained below normal levels after both RO and BV treatments while after BD it was reduced in central and peripheral regions (C: 38%±6%, p<0.0001; P: 59%±6%, p<0.0001). Thus, these results provided preliminary evidence that these treatments do not return normal levels of expression for all 7 proteins in all regions of V1. Instead, there was an intricate pattern of changes that did not clearly distinguish if a treatment shifted the overall pattern towards normal.

### Analyzing heterogeneity in protein expression and comparing among rearing conditions

We were surprised by the complexity in the pattern of protein changes and wondered if there was an abnormally high degree of heterogeneity in the protein expression. To assess heterogeneity, we calculated an index of dispersion (variance-to-mean ratio, VMR) for each protein and condition and used it to determine if the distributions were under-dispersed (VMR < 1), randomly-dispersed (VMR=1) or over-dispersed (VMR>1). For 5wk normal and RO conditions all proteins were under-dispersed with VMRs <0.3 (Figure 2). For the other conditions, the proteins were also under-dispersed except after MD. GluN2B was randomly-dispersed in the monocular region (Figure 2), and GABA_A_α1 was over-dispersed centrally after BD and in the monocular region after 4d BV. This analysis did not find increased heterogeneity that might explain the complexity.

**Figure 2.**
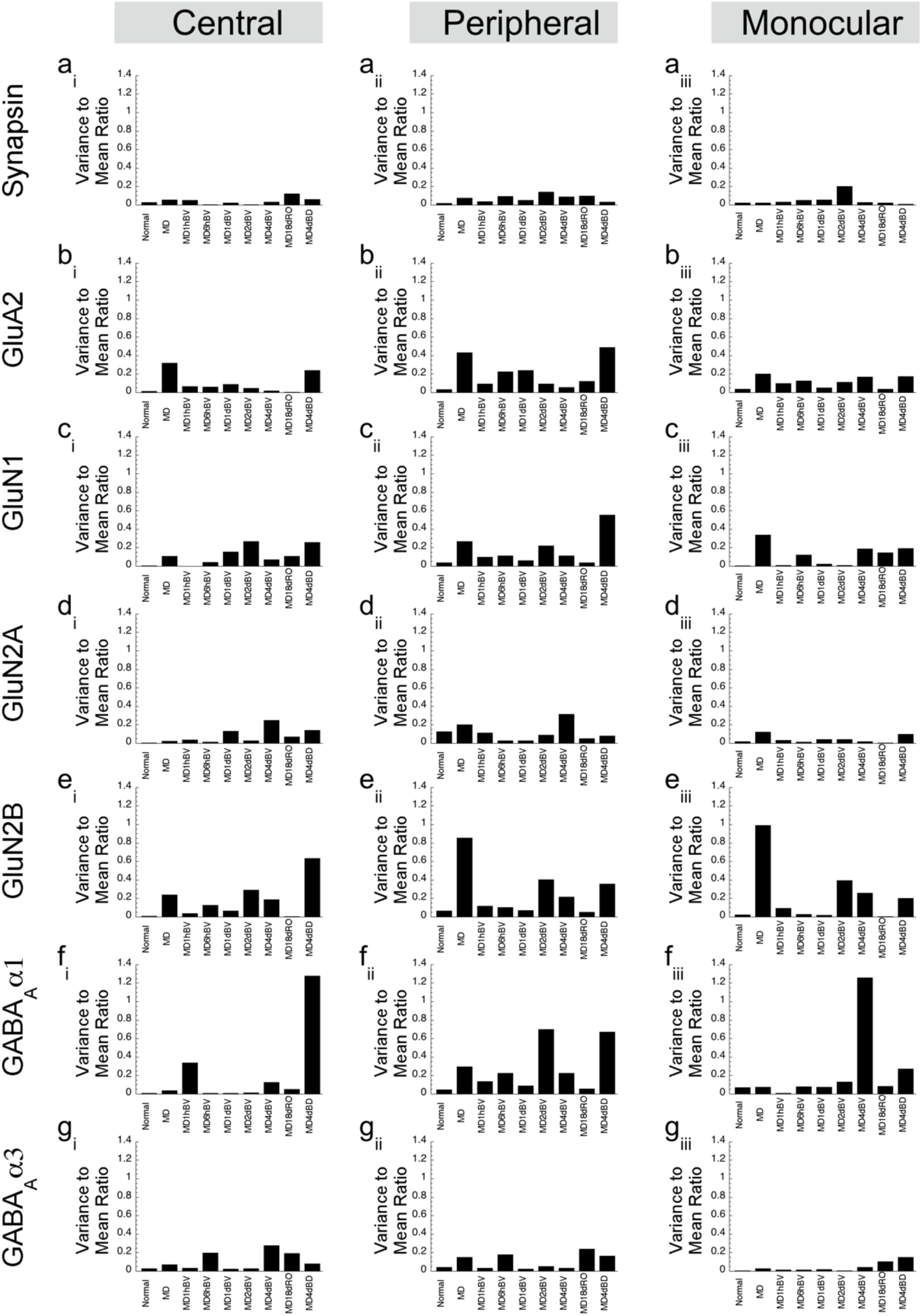
Glutamatergic and GABAergic receptor variance to mean ratios across conditions. Histograms depict the variance-to-mean ratio in each condition for individual proteins (rows) in each cortical area (columns). VMR >1 represent proteins that are highly-dispersed, VMR=1 are normally dispersed and VMR<1 are under-dispersed.

## Discussion

We quantified changes to a set of glutamatergic and GABAergic receptor subunits in V1 after either BD, RO or BV was given to promote recovery from MD. There were many changes, and none of the conditions restored normal expression to all of the different subunits. Furthermore, recovery often differed across the regions of V1 representing central, peripheral and monocular visual fields. RO was the only treatment that changed the expression of the pre-synaptic marker synapsin suggesting either more synapses or increased pre-synaptic functioning. BD appears to reduce both NMDA and AMPA receptor subunits but increased GABA_A_α1 that may result in a shift towards inhibition. In contrast, BV seemed to enhance glutamatergic subunits, especially NMDARs while also reducing GABA_A_Rs to cause a shift in favor of excitation. Since BV is the only treatment that promotes long-lasting visual recovery (Williams et al., 2015), the current findings suggest that enhanced excitatory drive is one of the mechanisms that supports those improvements in visual acuity.

## Conclusions

This study highlights changes to individual glutamatergic and GABAergic receptor subunits caused by BV, RO or BD used to promoted recovery from MD. Not surprisingly, the findings demonstrate the complex nature of the changes. There are hundreds of pre-synaptic and thousands of post-synaptic proteins that function as a balanced dynamic system to regulate neurotransmission and experience-dependent plasticity. Unravelling the synaptic proteins that support healthy versus abnormal functioning will need to use a systems approach and modern high-dimensional analyses to reveal the patterns of expression that are crucial for normal visual perception.

## Acknowledgements

We thank Kyle Hornby and Dr Brett Beston for assistance with data collection.

## Data Availability

The data used to support the findings of this study are available from the corresponding author upon request.

