## Supplementary Materials for "Recovery of glutamatergic and GABAergic protein expression in visual cortex after monocular deprivation"

| Weeks BV | Region | GluN1 | GluN2A | GluN2B | GABA$_A$α1 | GABA$_A$α3 | GluA2 | Synapsin |
|---|---|---|---|---|---|---|---|---|
| | C | 4 | 4 | 4 | 4 | 4 | 4 | 4 |
| 2 | P | 16 | 16 | 16 | 16 | 16 | 16 | 16 |
| | M | 4 | 4 | 4 | 4 | 4 | 4 | 4 |
| | C | 4 | 4 | 4 | 4 | 4 | 4 | 4 |
| 3 | P | 16 | 16 | 16 | 16 | 16 | 16 | 16 |
| | M | 4 | 4 | 4 | 4 | 4 | 4 | 4 |
| | C | 4 | 4 | 4 | 4 | 4 | 4 | 4 |
| 4 | P | 16 | 16 | 16 | 16 | 16 | 16 | 16 |
| | M | 4 | 4 | 4 | 4 | 4 | 4 | 4 |
| | C | 4 | 4 | 4 | 4 | 4 | 4 | 4 |
| 5 | P | 16 | 16 | 16 | 15 | 16 | 16 | 16 |
| | M | 4 | 4 | 4 | 4 | 4 | 4 | 4 |
| | C | 4 | 4 | 4 | 4 | 4 | 4 | 4 |
| 6 | P | 16 | 16 | 16 | 16 | 16 | 16 | 16 |
| | M | 4 | 4 | 4 | 4 | 4 | 4 | 4 |
| | C | 6 | 6 | 6 | 6 | 6 | 6 | 6 |
| 8 | P | 22 | 19 | 22 | 22 | 22 | 22 | 22 |
| | M | 2 | 2 | 2 | 2 | 2 | 2 | 2 |
| | C | 4 | 4 | 4 | 4 | 4 | 4 | 4 |
| 12 | P | 16 | 16 | 16 | 16 | 16 | 13 | 16 |
| | M | 4 | 4 | 4 | 4 | 4 | 4 | 4 |
| | C | 4 | 4 | 4 | 4 | 4 | 4 | 4 |
| 16 | P | 16 | 16 | 16 | 16 | 16 | 16 | 16 |
| | M | 4 | 4 | 4 | 4 | 4 | 4 | 4 |
| | C | 4 | 4 | 4 | 4 | 4 | 4 | 4 |
| 32 | P | 16 | 16 | 16 | 16 | 16 | 16 | 16 |
| | M | 4 | 4 | 4 | 4 | 4 | 4 | 4 |
| SUM | | 222 | 219 | 222 | 221 | 222 | 219 | 222 |

**Table 2-1. The number of Western blot measurements for each cortical region in Normal animals.** Rows summarize the number of runs from the Central (C), Peripheral (P), and Monocular (M) regions of V1 within each age of animal studied. The columns list each of the 7 proteins analyzed using Western blotting. Column sums detail the number of runs across ages and cortical areas.

| Weeks MD | Region | GluN1 | GluN2A | GluN2B | GABA$_A$α1 | GABA$_A$α3 | GluA2 | Synapsin |
|---|---|---|---|---|---|---|---|---|
| | C | 4 | 4 | 4 | 4 | 4 | 4 | 4 |
| 4 | P | 16 | 16 | 16 | 16 | 16 | 11 | 16 |
| | M | 4 | 4 | 4 | 4 | 4 | 2 | 4 |
| | C | 6 | 6 | 6 | 6 | 6 | 6 | 4 |
| 5 | P | 18 | 18 | 18 | 18 | 18 | 18 | 11 |
| | M | 4 | 4 | 4 | 4 | 4 | 4 | 4 |
| | C | 8 | 8 | 8 | 8 | 8 | 8 | 8 |
| 6 | P | 32 | 32 | 32 | 32 | 32 | 32 | 32 |
| | M | 6 | 6 | 6 | 6 | 6 | 6 | 6 |
| | C | 8 | 8 | 8 | 8 | 8 | 8 | 8 |
| 9 | P | 36 | 36 | 34 | 34 | 36 | 36 | 31 |
| | M | 8 | 8 | 8 | 6 | 8 | 8 | 6 |
| | C | 4 | 4 | 4 | 4 | 4 | 4 | 4 |
| 32 | P | 16 | 16 | 16 | 16 | 16 | 16 | 16 |
| | M | 2 | 2 | 2 | 2 | 2 | 2 | 2 |
| SUM | | 172 | 172 | 170 | 168 | 172 | 165 | 156 |

**Table 2-2. The number of Western blot measurements for each cortical region in MD animals.** Rows summarize the number of runs from the Central (C), Peripheral (P), and Monocular (M) regions of V1 within each age of animal studied. The columns list each of the 7 proteins analyzed using Western blotting. Column sums detail the number of runs across ages and cortical areas.

| | | Central | | | | | Peripheral | | | | | Monocular | | | | |
|---|---|---|---|---|---|---|---|---|---|---|---|---|---|---|---|---|
| | | Vs 5wk Normal | | vs. 5wk MD | | Curve Fit to BV Data | Vs 5wk Normal | | vs. 5wk MD | | Curve Fit to BV Data | Vs 5wk Normal | | vs. 5wk MD | | Curve Fit to BV Data |
| | | Significance | p-value | Significance | p-value | | Significance | p-value | Significance | p-value | | Significance | p-value | Significance | p-value | |
| Synapsin | MD | n.s. | 0.2361 | | | | n.s. | 0.2113 | | | | n.s. | 0.4861 | | | |
| | RO | *** | 0.0000 | *** | 0.0000 | | *** | 0.0000 | *** | 0.0000 | | *** | 0.0000 | *** | 0.0000 | |
| | BD | n.s. | 0.1483 | * | 0.0123 | | n.s. | 0.2252 | n.s. | 0.0526 | | *** | 0.0001 | *** | 0.0001 | |
| | 1hr BV | n.s. | 0.1438 | * | 0.0107 | | *** | 0.0000 | ** | 0.0026 | | *** | 0.0000 | *** | 0.0000 | |
| | 6hr BV | *** | 0.0000 | *** | 0.0000 | | * | 0.0152 | n.s. | 0.1051 | | n.s. | 0.2726 | n.s. | 0.2829 | |
| | 1d BV | *** | 0.0009 | *** | 0.0000 | | *** | 0.0000 | *** | 0.0001 | | ** | 0.0095 | ** | 0.0095 | |
| | 2d BV | *** | 0.0000 | *** | 0.0000 | | n.s. | 0.1742 | n.s. | 0.1002 | | n.s. | 0.1263 | n.s. | 0.1215 | |
| | 4d BV | ** | 0.0029 | *** | 0.0000 | | n.s. | 0.1474 | n.s. | 0.0668 | | n.s. | 0.3185 | n.s. | 0.3037 | |
| GluA2 | MD | n.s. | 0.2338 | | | | ** | 0.0085 | | | | n.s. | 0.3516 | | | |
| | RO | *** | 0.0000 | *** | 0.0000 | | n.s. | 0.0691 | *** | 0.0000 | | *** | 0.0000 | *** | 0.0000 | |
| | BD | *** | 0.0000 | *** | 0.0000 | | *** | 0.0001 | n.s. | 0.1110 | | n.s. | 0.2511 | n.s. | 0.1042 | |
| | 1hr BV | *** | 0.0000 | *** | 0.0006 | | *** | 0.0000 | *** | 0.0000 | | *** | 0.0000 | *** | 0.0000 | |
| | 6hr BV | *** | 0.0007 | n.s. | 0.4871 | | n.s. | 0.1370 | * | 0.0253 | | * | 0.0418 | n.s. | 0.0760 | |
| | 1d BV | *** | 0.0003 | * | 0.0106 | | *** | 0.0000 | * | 0.0405 | | n.s. | 0.0912 | n.s. | 0.2506 | |
| | 2d BV | *** | 0.0000 | *** | 0.0000 | | *** | 0.0000 | *** | 0.0000 | | * | 0.0106 | *** | 0.0006 | |
| | 4d BV | n.s. | 0.2257 | ** | 0.0051 | | *** | 0.0000 | n.s. | 0.1718 | | *** | 0.0006 | *** | 0.0000 | |
| GluN1 | MD | *** | 0.0000 | | | | ** | 0.0038 | | | | n.s. | 0.1573 | | | |
| | RO | ** | 0.0041 | * | 0.0104 | | *** | 0.0000 | *** | 0.0002 | | ** | 0.0019 | *** | 0.0000 | |
| | BD | n.s. | 0.1908 | * | 0.0123 | | n.s. | 0.1753 | n.s. | 0.1933 | | n.s. | 0.4286 | * | 0.0292 | |
| | 1hr BV | *** | 0.0000 | n.s. | 0.0856 | | *** | 0.0000 | n.s. | 0.2717 | | ** | 0.0014 | *** | 0.0000 | |
| | 6hr BV | *** | 0.0000 | *** | 0.0001 | y = 297.43-239.58*exp(-x/11.61) | n.s. | 0.4143 | *** | 0.0002 | | n.s. | 0.3155 | n.s. | 0.0706 | y = 62.45+51.87*exp(-x/0.79) |
| | 1d BV | * | 0.0294 | * | 0.0144 | df=25 | *** | 0.0000 | * | 0.0366 | | *** | 0.0001 | *** | 0.0000 | df=24 |
| | 2d BV | n.s. | 0.3664 | ** | 0.0039 | $R^2$=0.618 | *** | 0.0000 | n.s. | 0.1493 | | *** | 0.0000 | *** | 0.0000 | $R^2$=0.361 |
| | 4d BV | ** | 0.0044 | *** | 0.0000 | p<0.0001 | n.s. | 0.2547 | *** | 0.0000 | | * | 0.0214 | *** | 0.0000 | p=0.0012 |
| GluN2A | MD | *** | 0.0000 | | | | *** | 0.0000 | | | | * | 0.0220 | | | |
| | RO | n.s. | 0.1367 | *** | 0.0007 | | *** | 0.0000 | * | 0.0142 | | *** | 0.0000 | *** | 0.0000 | |
| | BD | *** | 0.0000 | n.s. | 0.0923 | | *** | 0.0000 | *** | 0.0000 | | *** | 0.0000 | *** | 0.0000 | |
| | 1hr BV | *** | 0.0001 | * | 0.0220 | | *** | 0.0000 | *** | 0.0007 | | *** | 0.0005 | n.s. | 0.2953 | |
| | 6hr BV | *** | 0.0001 | *** | 0.0000 | y = 671.85-601.29*exp(-x/39.72) | *** | 0.0000 | *** | 0.0000 | | *** | 0.0000 | n.s. | 0.2208 | y = 52.37+(77.48-52.37)/(1+(x/1.06)^14.48) |
| | 1d BV | n.s. | 0.2788 | ** | 0.0065 | df=23 | *** | 0.0000 | *** | 0.0000 | | *** | 0.0000 | n.s. | 0.1832 | df=24 |
| | 2d BV | n.s. | 0.2635 | *** | 0.0000 | $R^2$=0.528 | *** | 0.0000 | n.s. | 0.4093 | | *** | 0.0000 | *** | 0.0000 | $R^2$=0.422 |
| | 4d BV | n.s. | 0.1022 | *** | 0.0002 | p<0.0001 | *** | 0.0003 | n.s. | 0.0957 | | *** | 0.0000 | *** | 0.0000 | p=0.0003 |
| GluN2B | MD | n.s. | 0.4587 | | | | n.s. | 0.1907 | | | | n.s. | 0.0606 | | | |
| | RO | *** | 0.0000 | *** | 0.0000 | | *** | 0.0000 | *** | 0.0000 | | *** | 0.0000 | *** | 0.0000 | |
| | BD | ** | 0.0085 | ** | 0.0013 | | *** | 0.0000 | *** | 0.0000 | | * | 0.0354 | *** | 0.0000 | |
| | 1hr BV | n.s. | 0.4438 | n.s. | 0.3090 | | ** | 0.0059 | *** | 0.0000 | | * | 0.0374 | *** | 0.0000 | |
| | 6hr BV | n.s. | 0.0574 | * | 0.0384 | | ** | 0.0033 | n.s. | 0.3986 | | * | 0.0172 | *** | 0.0000 | y = 80.23+137.58*exp(-x/0.04) |
| | 1d BV | n.s. | 0.1647 | n.s. | 0.0745 | | n.s. | 0.1926 | ** | 0.0050 | | n.s. | 0.0893 | *** | 0.0000 | df=24 |
| | 2d BV | n.s. | 0.3470 | n.s. | 0.3570 | | n.s. | 0.0574 | *** | 0.0004 | | n.s. | 0.1156 | *** | 0.0000 | $R^2$= 0.396 |
| | 4d BV | n.s. | 0.4151 | n.s. | 0.4538 | | ** | 0.0024 | *** | 0.0000 | | *** | 0.0007 | *** | 0.0000 | p= 0.0006 |
| $GABA_A\alpha1$ | MD | *** | 0.0000 | | | | *** | 0.0007 | | | | *** | 0.0000 | | | |
| | RO | * | 0.0239 | *** | 0.0000 | | *** | 0.0008 | *** | 0.0000 | | ** | 0.0026 | n.s. | 0.1754 | |
| | BD | n.s. | 0.0977 | n.s. | 0.1914 | | *** | 0.0000 | *** | 0.0000 | | *** | 0.0000 | *** | 0.0000 | |
| | 1hr BV | n.s. | 0.1903 | n.s. | 0.2725 | | * | 0.0357 | *** | 0.0000 | | *** | 0.0002 | *** | 0.0000 | |
| | 6hr BV | n.s. | 0.2956 | *** | 0.0000 | y = 57.31+77.01*exp(-x/0.47) | *** | 0.0000 | ** | 0.0070 | | *** | 0.0000 | *** | 0.0000 | |
| | 1d BV | *** | 0.0000 | *** | 0.0000 | df=24 | ** | 0.0022 | *** | 0.0000 | | n.s. | 0.2356 | ** | 0.0011 | |
| | 2d BV | *** | 0.0006 | *** | 0.0000 | $R^2$=0.636 | n.s. | 0.0562 | *** | 0.0000 | | *** | 0.0000 | * | 0.0179 | |
| | 4d BV | *** | 0.0000 | *** | 0.0000 | p<0.0001 | *** | 0.0000 | *** | 0.0000 | | *** | 0.0000 | *** | 0.0000 | |
| $GABA_A\alpha3$ | MD | *** | 0.0010 | | | | *** | 0.0001 | | | | *** | 0.0000 | | | |
| | RO | n.s. | 0.0614 | n.s. | 0.3711 | | ** | 0.0015 | n.s. | 0.4668 | | ** | 0.0094 | n.s. | 0.0883 | |
| | BD | *** | 0.0000 | *** | 0.0000 | | *** | 0.0000 | * | 0.0141 | | n.s. | 0.4510 | ** | 0.0040 | |
| | 1hr BV | *** | 0.0001 | n.s. | 0.2468 | | *** | 0.0000 | *** | 0.0000 | | *** | 0.0000 | ** | 0.0031 | |
| | 6hr BV | n.s. | 0.0524 | n.s. | 0.4068 | y = 40.25+33.81*exp(-x/2.87) | *** | 0.0000 | n.s. | 0.0917 | y = 47.66+20.83*exp(-x/0.51) | n.s. | 0.2084 | *** | 0.0000 | |
| | 1d BV | *** | 0.0000 | *** | 0.0001 | df=25 | *** | 0.0000 | *** | 0.0000 | df=91 | *** | 0.0000 | *** | 0.0007 | |
| | 2d BV | *** | 0.0000 | ** | 0.0047 | $R^2$=0.211 | *** | 0.0000 | *** | 0.0000 | $R^2$=0.163 | *** | 0.0000 | n.s. | 0.4088 | |
| | 4d BV | *** | 0.0003 | * | 0.0229 | p=0.0159 | *** | 0.0000 | *** | 0.0000 | <0.0001 | *** | 0.0000 | n.s. | 0.1572 | |

**Table 3-1. Table of p-values comparing protein expression in each treatment condition against 5wk Normal animals and 5wk MD animals.** P-values are presented for each cortical area (columns) and protein (rows). Cortical areas are broken up into comparisons against normal (left) and MD (right). Asterisk color coding matches Figure 3. When a curve fit was applied, the equation, degrees of freedom (df), $R^2$ value and exact p-value are listed.
